# Surfactant protein A promotes atherosclerosis through mediating macrophage foam cell formation

**DOI:** 10.1101/2023.03.23.533959

**Authors:** Skylar D. King, Dunpeng Cai, Mikayla M. Fraunfelder, Shi-You Chen

## Abstract

**BACKGROUND:** Atherosclerosis is a progressive inflammatory disease where macrophage foam cells play a central role in the pathogenesis. Surfactant protein A (SPA) is a lipid-associating protein involved with regulating macrophage function in various inflammatory diseases. However, the role of SPA in atherosclerosis and macrophage foam cell formation has not been investigated.

**METHODS:** Primary resident peritoneal macrophages were extracted from wildtype (WT) and SPA deficient (SPA^-/-^) mice to determine the functional effects of SPA in macrophage foam cell formation. SPA expression was assessed in healthy vessels and atherosclerotic aortic tissue from the human coronary artery and WT or apolipoprotein e-deficient (ApoE^-/-^) mice brachiocephalic arteries fed high fat diets (HFD) for 4 weeks. Hypercholesteremic WT and SPA^-/-^ mice fed a HFD for 6 weeks were investigated for atherosclerotic lesions *in vivo*.

**RESULTS:** *In vitro* experiments revealed that global SPA deficiency reduced intracellular cholesterol accumulation and macrophage foam cell formation. Mechanistically, SPA^-/-^ dramatically decreased CD36 cellular and mRNA expression. SPA expression was increased in atherosclerotic lesions in humans and ApoE^-/-^ mice. *In vivo* SPA deficiency attenuated atherosclerosis and reduced the number of lesion-associated macrophage foam cells.

**CONCLUSIONS:** Our results elucidate that SPA is a novel factor for atherosclerosis development. SPA enhances macrophage foam cell formation and atherosclerosis through increasing scavenger receptor cluster of differentiation antigen 36 (CD36) expression.

## INTRODUCTION

Cardiovascular disease is the most common cause of mortality and morbidity world-wide^1,2^. Atherosclerosis (AS) is a progressive disease characterized by persistent inflammation and excessive cholesterol deposition within the artery wall and is the fundamental pathological process underlying coronary artery disease and stroke^3^. Macrophages play a critical role in AS development because of their ability to become lipid laden foam cells^4^. Macrophage-derived foam cell formation begins with monocyte recruitment from circulation into the subendothelial space of the affected arterial wall, leading to lipid metabolism imbalance and maladaptive immune responses^4^. The subendothelial space is a cytokine-rich microenvironment where monocytes differentiate into macrophages and express proteins that mediate the retention of cholesterol in cells and the uptake of modified lipoproteins, such as oxidized low-density lipoprotein (OxLDL)^5^. Understanding the mechanisms whereby macrophages accumulate LDL cholesterol is essential to the understanding of early AS development and progression.

The accumulation of lipid-laden foam cells within the vessel plays a crucial role in the initiation and progression of AS^5^. The transformation of macrophages into foam cells is controlled by the lipid uptake of modified lipoproteins regulated by scavenger receptors contributing to fatty-streak formation, specifically, scavenger receptor A (SRA) and cluster of differentiation antigen 36 (CD36). CD36 is responsible for an estimated 75% OxLDL uptake^6^.

Surfactant protein A (SPA) is a large glycoprotein in the collectin family of collagen-like lectins^7^. SPA has been studied in pulmonary immune defense and alveolar macrophage lipid homeostasis and is present in extrapulmonary tissues^7,8^. SPA has been implicated in lung-specific host defense because of its interactions with a diversity of pathogens and due to stimulating chemotaxis, phagocytosis, secretion of colony-stimulating factors and reactive oxygen intermediates by alveolar macrophages^7,9^–^12^. In view of the critical role of SPA in macrophages and the essential functions of macrophages in AS development, we speculated that SPA may play a role in the development or progression of AS. Together, these findings highlight the importance of SPA in the regulation of immune and inflammatory responses in health and diseases. However, the role of SPA in macrophage foam cell formation (MFCF) and AS has not yet been investigated. Given that AS is associated with dyslipidemia and inflammation, we hypothesized that SPA functions to enhance MFCF formation resulting in increased AS. This study aimed to provide evidence to elucidate the mechanism of SPA in MFCF and AS. Thus, we explored for the first time the involvement of SPA in MFCF and AS. Our results reveal that SPA deficiency *in vitro* reduces the accumulation of OxLDL-induced cholesterol esters and free cholesterol and the production of OxLDL-induced MFCF. We further found that SPA deficiency inhibits OxLDL induced CD36 cellular and mRNA expression. I*n vivo* studies show that SPA is expressed in atherosclerotic lesions and its deficiency attenuates AS lesion formation. Together, these results reveal that SPA is a novel regulator promoting the key initial process of AS (fatty streak formation).

## MATERIALS AND METHODS

All materials used are included in the **Major Resources Table**. Detailed methods are in the **Supplementary Materials**.

### Data Availability

The authors declare that all of the relevant data supporting the findings of this study are available from the corresponding author upon reasonable request.

### Experimental animals

SPA deficient (SPA^-/-^) (*Sftpa1^tm1Kor^*/J, strain number: 004964)^13^, ApoE^-/-^ (B6.129-*Apoe^tm1Unc^*/J, strain number: 002052) and wild-type (C57BL6/J, strain number: 000664) mice were purchased from the Jackson Laboratory. The SPA^-/-^ mice have been crossed with C57BL6/J for more than 10 generations. Genotyping primers are included in the Major Resources Table. All animals used in this study were housed under conventional conditions in the animal care facilities and received humane care in compliance with the Principles of Laboratory Animal Care formulated by the National Society of Medical Research and the Guide for the Care and Use of Laboratory Animals. All experimental procedures were approved by the Institutional Animal Care and Use Committee of the University of Missouri, IACUC Project number 9736, and followed the recommendations in the ARRIVE guidelines.

### Macrophage cell culture and treatment

Resident peritoneal macrophages were isolated from 6 to 8-week-old WT or SPA^-/-^ male or female mice as previously described^14^. Peritoneal fluid was collected in sterile PBS and centrifuged at 400 *g* for 10 minutes. The cells were washed with PBS and plated. One hour after incubation, nonadherent cells were removed. Adhered macrophages were cultured in DMEM medium supplemented with 10% FBS, 2 mM L-glutamine, 100U/mL penicillin and 100 μg/mL streptomycin at 37°C in a humidified atmosphere with 5% CO^2^ and used for future experiments. For experiments involving treatment, cells were immediately used from the peritoneal cavity, allowed to adhere to coverslip in supplemented DMEM media then changed to (1) serum-free DMEM (Control) or (2) serum-free DMEM containing 50 μg/ml OxLDL, for 24 hours (h).

### Human coronary artery tissue protein extraction and staining

The protocol for using human tissue samples was approved by the Institutional Review Board (IRB) at the University of Missouri-Columbia, IRB Project number 2026026. All experiments conducted with human specimens were performed in accordance with the relevant regulations and guidelines. Formalin fixed paraffin-embedded (FFPE) segments of human coronary arteries with normal or atherosclerotic arteries were obtained from autopsy materials. SPA expression was detected from proteins extracted from the human coronary artery FFPE tissues^18^. For immunofluorescent staining, 5 μm sections were cut, dewaxed, rehydrated, and boiled for 3 minutes in 0.05% Tween 20 citrate buffer (10mmol/L, pH6) for antigen retrieval and then blocked in 10% normal goat serum for 30 min. Blocked sections were incubated overnight at 4°C with primary antibodies SPA (1:100) or CD68 (1:200) followed by incubation with goat anti-mouse IgG Alexafluor 488 or goat antirabbit IgG (H+L) Alexafluor 564. Coverslips were applied over the tissue sections with Prolong Gold antifade medium containing DAPI. Images were captured using a Keyence microscope and were processed using ImageJ.

### Statistical analysis

Each experiment was repeated at least 3 times with at least 2 technical repeats. All data represent independent data points within all technical repeats. Values are expressed as the mean with standard deviation. Data values were analyzed by comparing experimental values to control values by analyzing for Gaussian distribution using D’ Agostino & Pearson and Shapiro-Wilk normality tests (alpha=0.05, p<0.05). If normality was passed, parametric statistical test, unpaired t-test with Welch’s correction or Ordinary one-way ANOVA followed by Tukey’s multiple comparisons test, was performed. If normality was not passed, or for groups with n less than 6, data was analyzed using nonparametric statistical tests, unpaired t-test Mann-Whitney test or ANOVA Kruskal-Wallis multiple comparisons test. Statistical analysis was conducted using GraphPad Prism 8 software, and differences were considered statistically significant when nominal p<0.05 or adjusted p<0.05 in case of multiple testing. Correction for multiple testing across the entire body of experiments was not performed because both *in vitro* and *in vivo* experiments were performed and various approaches were used in this study. All p-values and the corresponding statistical tests are provided in the **Online Data Supplement Table 1**.

## RESULTS

### SPA deficiency attenuates intracellular cholesterol accumulation and OxLDL-induced MFCF

Foam cell formation is the hallmark of the initiation and development of AS^5^. The major core components of low- and high-density lipoproteins are highly nonpolar cholesteryl ester lipids^21^. BODIPY binds specifically to cellular lipid droplets and the cholesteryl ester content of cells can be monitored by the emitted fluorescence^22,23^. To test if SPA is involved in OxLDL-induced MFCF, we extracted peritoneal MΦ from WT and SPA^-/-^ mice and evaluated the effects of cholesterol, OxLDL uptake and foam cell formation after 24 h of OxLDL treatment. We first determined SPA expression via IF staining in WT or SPA^-/-^ MΦ and found that SPA staining was not observed in the MΦs extracted from SPA^-/-^ mice, confirming the knockout efficiency *Sftpa1^tm1Kor^*/J mice (**Figure 1A**)^13^. Next, we evaluated the effect of cholesterol esters and intracellular free cholesterol accumulation in MΦ collected from WT or SPA^-/-^ mice upon exposure to OxLDL for 24 h via BODIPY or Filipin III staining (**Figure 1, B & D**). Consistent with what others have found, OxLDL induced intracellular cholesterol accumulation relative to the control^24,25^. We further found that the amounts of cholesterol esters and free intracellular cholesterol were significantly decreased in SPA^-/-^ MΦs relative to the WT cells treated with OxLDL (**Figure 1, C & E**). Additionally, we incubated with Dil-conjugated OxLDL for 24 h and compared the amount of internalized Dil-conjugated OxLDL (Dil-OxLDL) between WT or SPA^-/-^ cells (**Figure 1F**), and found that SPA deficiency significantly decreased the amount of Dil-OxLDL uptake and consequently the percentage of foam cell formation relative to WT MΦs (**Figure 1, G-H**). Together, these results reveal that SPA deficiency reduced intracellular cholesterol esters and free cholesterol, Dil-OxLDL uptake and MFCF *in vitro*, suggesting that the presence of SPA is essential for macrophage foam cell formation.

**Figure 1.**
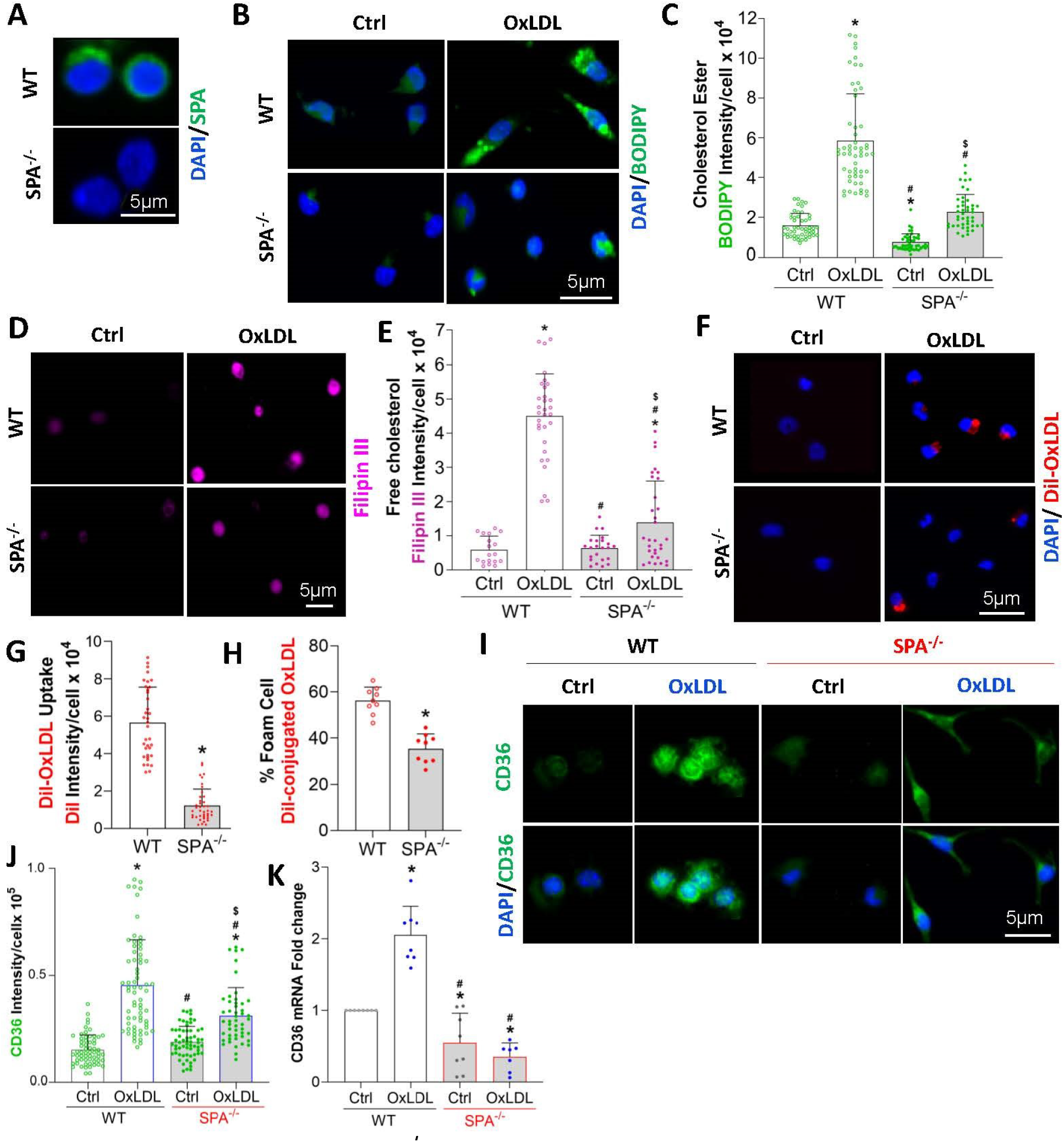
SPA deficiency (SPA^-/-^) diminishes macrophage (MΦ) foam cell formation through inhibiting CD36 expression *in vitro*. Primary resident peritoneal MΦs isolated from wild-type (WT) or SPA^-/-^ mice were treated with vehicle (Ctrl) or OxLDL (50 μg) for 24 hours. **(A)** Representative immunofluorescence images of MΦs stained with SPA antibody (green) confirming SPA deletion in SPA^-/-^ MΦs. Nuclei stained with DAPI (blue). **(B)** Representative images of neutral lipids (cholesterol ester) accumulation as detected by BODIPY staining (green). Nuclei stained with DAPI (blue). **(C)** Quantification of the BODIPY fluorescence intensity per MΦ. Each dot represents the intensity of a single MΦ. * p<0.0003 vs. Ctrl WT MΦs; # p<0.0001 vs. OxLDL-treated WT MΦs; $ p<0.0001 vs. Ctrl SPA^-/-^ MΦs. Krustal-Wallis test followed by Dunn’s multiple comparisons test; WT MΦs; Ctrl n=49, OxLDL n=55; SPA^-/-^ MΦs; Ctrl n=45, OxLDL n=45. **(D)** Representative images of intracellular free cholesterol content as detected by Filipin III staining, (magenta). **(E)** Quantification of free cholesterol content by measuring the Filipin III staining intensity. Each dot represents the intensity of a single MΦ. * p<0.034 vs. Ctrl WT MΦs; # p<0.0001 vs. OxLDL-treated WT MΦs; $ p=0.031 vs. Ctrl SPA^-/-^ MΦs). One-way ANOVA with Tukey’s multiple comparisons test, WT MΦs; Ctrl n=18, OxLDL n=31; SPA^-/-^ MΦs; Ctrl n=23, OxLDL n=31. **(F)** Representative images of OxLDL uptake by WT and SPA^-/-^ MΦs as displayed by the presence of Dil-OxLDL (red). Nuclei stained with DAPI (blue). **(G)** Quantification of OxLDL uptake by the fluorescence intensity of Dil-conjugated OxLDL. Each dot represents the intensity of a single cell. * p<0.0001 vs. WT MΦs; Mann Whitney test; WT MΦs; n=36; SPA^-/-^ MΦs; n=40. **(H)** Quantification of foam cell formation as % cells containing Dil-conjugated OxLDL relative to the total number of MΦs. * p=0.0001 vs. WT MΦs; Unpaired two-tailed t test with Welch’s correction, n=9. **(I)** Representative images of CD36 antibody staining (green). Nuclei stained with DAPI (blue) **(J)** Quantification of CD36 expression by measuring green fluorescent intensity. Each dot represents the intensity of a single vs. WT MΦs. * p<0.0001 vs. Ctrl WT MΦs; # p<0.046 vs. OxLDL-treated WT MΦs; $ p<0.0001 vs. Ctrl SPA^-/-^ MΦs; Kruskal-Wallis test followed by Dunn’s multiple comparisons test, WT MΦs; Ctrl n=60, OxLDL n=67; SPA^-/-^ MΦs; Ctrl n=62, OxLDL n=88. **(K)** qRT-PCR analysis of CD36 mRNA expression in MΦs. The fold change of CD36 expression was normalized to CYP expression and relative to Ctrl WT MΦs (set as 1). * and Indicates significance * p value ≥0.032 vs. Ctrl WT MΦs; # p<0.0001 vs. OxLDL-treated WT MΦs. Ordinary one-way ANOVA followed by Tukey’s multiple comparisons test, n=7-8. For all data represented, at least 3 independent experiments were conducted with at least 2 technical repeats, n indicates the quantification of data within all experiments, values are not averaged.

### SPA deficiency reduces the expression of OxLDL scavenger receptor CD36

The formation of foam cells is associated with increased lipid influx. OxLDL uptake is dependent on scavenger receptors^6^. Studies with mice lacking SRA and CD36 identified CD36 as the major OxLDL receptor required for the formation of foam cells^13,26–28^. To determine whether the decreased OxLDL uptake in SPA^-/-^ MΦs is because of altered intracellular expression of CD36, we analyzed the cellular expression of CD36 in MΦ extracted from WT or SPA^-/-^ mice upon treatment with vehicle or OxLDL for 24 h. We observed increased intracellular staining of CD36 in WT MΦs treated with OxLDL relative to the nontreated controls, which is consistent with previous findings^29^. However, in the SPA^-/-^ MΦs, we detected significantly decreased CD36 intracellular expression upon OxLDL treatment compared to vehicle-treated SPA^-/-^ MΦs or WT cells (**Figure 1, I & J**). Together, these results suggest that SPA deficiency affects CD36 cellular expression, which could be the primary reason for the decreased MFCF observed in SPA deficient MΦs.

To determine whether SPA deficiency also affected the transcription of CD36, we investigated CD36 mRNA expression in MΦ extracted from WT and SPA^/-^ mice upon vehicle or OxLDL treatment for 24 h. We found that WT cells treated with OxLDL increased the mRNA expression of CD36 relative to the WT Control (**Figure 1K**). SPA deficiency inhibited the mRNA expression of CD36 relative to the WT control and blocked the increased expression induced by OxLDL in WT MΦs (**Figure 1K**). Together, these data further support that SPA affects MFCF through the regulation of scavenger receptor CD36, which indicates that the decreased macrophage transformation into foam cells observed in SPA^-/-^ cells is, at least partially, through decreasing OxLDL uptake.

### SPA is upregulated in macrophages of atherosclerotic lesions

To analyze the role of SPA in the initiation and progression of AS, we detected SPA protein levels in human coronary arteries with or without AS lesions using Western blot analyses of proteins extracted from formalin fixed paraffin-embedded tissues (**Figure 2A**). SPA protein expression was observably increased in the coronary artery tissues with AS (**Figure 2B**). Next, we aimed to detect SPA expression in human vessels, thus, we conducted immunofluorescent (IF) staining of human coronary arteries with the SPA antibody. SPA staining was undetectable in the control arteries, but was significantly increased in the diseased vessels (**Figure 2C**). Colocalization analysis with CD68 revealed that SPA colocalized with MΦs within the AS lesions (**Figure 2C**, white arrow). These results suggest a potential role for SPA in the development of atherosclerotic plaques.

**Figure 2.**
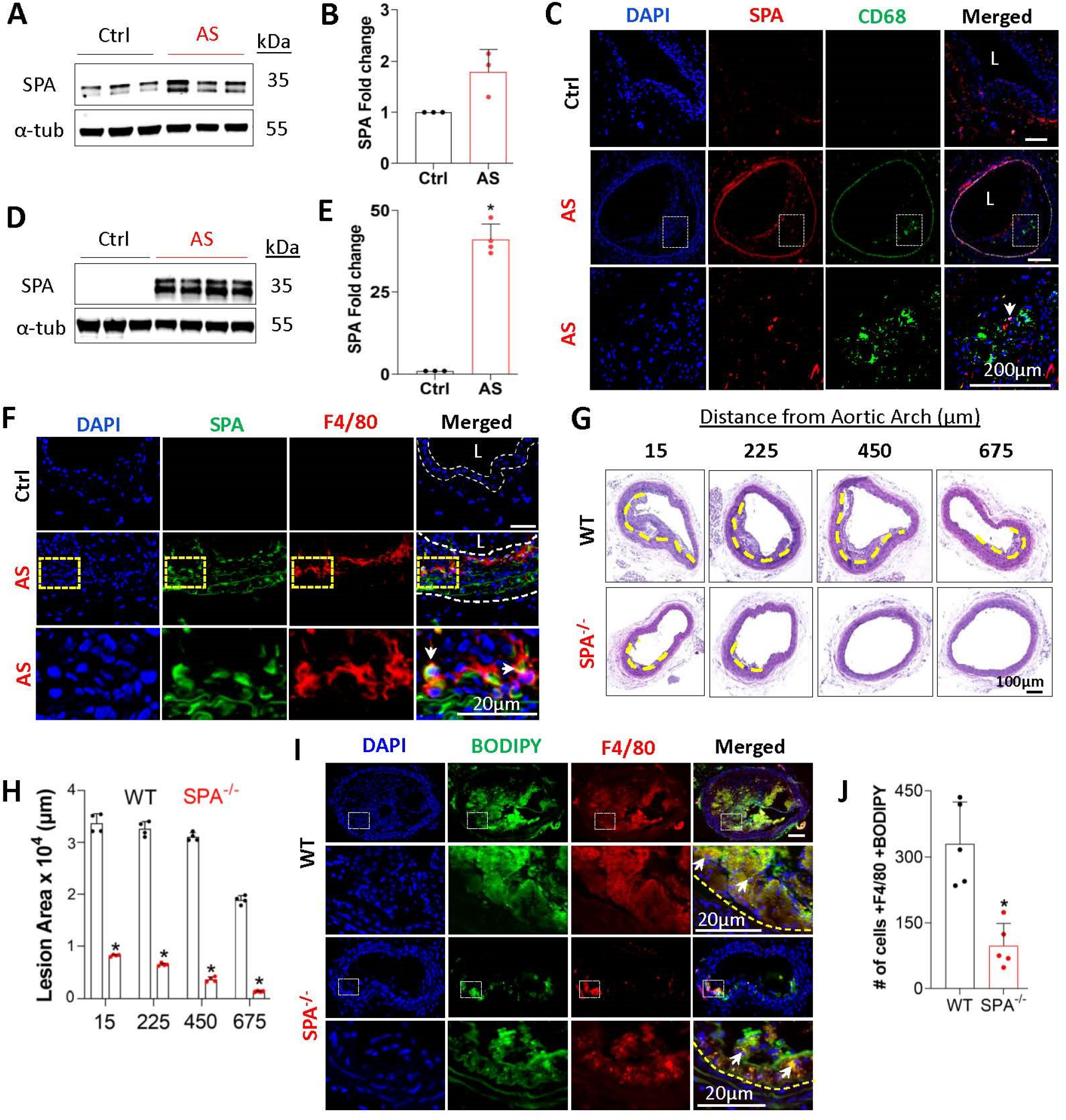
SPA deficiency attenuates atherosclerosis and macrophage foam cell formation *in vivo*. SPA expression was increased in coronary artery atherosclerotic lesion (AS) in human patients, as detected by Western blot **(A)** and normalized to alpha-Tubulin (α-Tub) **(B)** p=0.100 vs. health control. Nonparametric two-tailed Mann-Whitney test, n=3. **(C)** Coimmunostaining of SPA (red) with macrophage marker CD68 (green) nuclei stained with DAPI (blue) in human coronary artery atherosclerotic lesions. SPA expression was induced in brachiocephalic artery (BCA) of ApoE^-/-^ mice fed high fat diet for 4 weeks (AS), as detected by Western blot **(D)** and normalized to α-Tub **(E)** *p=0.029 vs. wild-type mice fed a high fat diet for 4 weeks (Ctrl). Nonparametric two-tailed Mann-Whitney test, n=3-4. **(F)** SPA was expressed in F4/80+ macrophages in BCA atherosclerotic lesions as detected by co-immunofluorescent staining. White dashed line outlines the lesion area. L: Lumen. The images in the yellow boxes are enlarged in the lower part of the panel. White arrows indicate colocalization of SPA with F4/80. **(G)** WT and SPA^-/-^ mice were injected with PCSK9-AAV followed by high fat diet feeding for 6 weeks. Atherosclerotic lesions in BCA sections were observed by H&E staining. Dashed yellow lines: boundary of the developing atherosclerotic lesions. **(H)** Quantification of the total plaque areas in serial sections from WT or SPA^-/-^ mice. (n=4). *p=0.029 vs. WT in each individual location. Nonparametric two-tailed Mann-Whitney test, n=4. **(I)** Representative images of foamy macrophages within mouse atherosclerotic lesions in the BCA as stained with BODIPY (green), F4/80 (red) and nuclei stained with DAPI (blue). The regions in dashed white squares are enlarged in the lower part of the panel for WT and SPA^-/-^, respectively. Yellow dashed line indicates the boundary of the intima. White arrows: macrophage foam cells. **(J)** Quantification of macrophage foam cells (BODIPY+F4/80+) relative to the total F4/80+ cells in atherosclerotic lesions. Each dot represents the total quantification from one animal. *p=0.008 vs. WT. Nonparametric two-tailed Mann-Whitney test, n=5.

In order to test if SPA is induced in the AS lesions of an animal model, we fed WT and ApoE^-/-^ mice a HFD for 4 weeks to initiate early AS development. BCAs were then collected and proteins extracted for Western blot analyses of SPA expression **(Figure 2D)**. Although control BCA exhibited a very low level of expression, SPA protein was significantly increased in the BCA of ApoE^-/-^ AS mice (**Figure 2E**). To determine the cellular localization of SPA within the AS lesions, we conducted IF staining with SPA antibody in the BCA. WT animals fed HFD had no visible atherosclerotic lesions or SPA expression (**Figure 2F**). Interestingly, in the mouse AS lesions, SPA expression was predominantly located within the media and the intima (**Figure 2F**). Consistent with our observations in the human AS lesion, SPA was also expressed in MΦs within mouse AS lesions as shown by its colocalization with F4/80 (**Figure 2F**, white arrow). Collectively, these results indicate that SPA is dramatically increased in MΦs within AS lesions, suggesting a potential role for SPA in macrophages and AS development.

### SPA deficiency attenuates the development of atherosclerosis and reduces macrophage-derived foam cells *in vivo*

To examine the potential role of SPA in AS *in vivo*, WT and SPA^-/-^ mice were administered with adenovirus-associated virus overexpressing PCSK9 (AAV-PCSK9) by tail vein injection to induce hypercholesterolemia^30^. Age and sex-matched animals were then fed a HFD for 6 weeks to observe early AS development. Quantification of the lesion burden by serial cross-sectional analyses of the BCA via H&E revealed that SPA deficiency reduced the AS lesion areas (**Figure 2, G-H**), indicating that loss of SPA inhibits atherosclerotic lesion progression.

To determine the effect of SPA on the accumulation of macrophage-derived foam cells within AS plaques, we determined whether SPA deficiency altered the number of foamy MΦs (F4/80+ MΦs containing lipid droplets, BODIPY^19^) within BCA sections (**Figure 2I**). Indeed, SPA deficiency significantly reduced the number of lesion associated F4/80+ MΦ foam cells (**Figure 2J**). These results demonstrate that SPA deficiency significantly reduces the number of MΦs that have been converted into foam cells, which is likely responsible for the reduced AS lesion formation in the BCA observed in SPA^-/-^ mice.

## DISCUSSION

In this study, we used primary mouse resident peritoneal macrophages from WT and SPA^-/-^ mice for *in vitro* studies and the hyperlipidemic atherosclerotic mouse model to investigate the impact of SPA deficiency on the formation of macrophage-derived foam cells and AS development *in vivo*. The inflammatory response of macrophages to invading pathogenic lipoproteins in the arterial wall and the formation of macrophage foam cells in the intima is a hallmark of early-stage atherosclerotic lesions^3,31^. Our *in vitro* studies revealed that SPA deficiency inhibited intracellular cholesterol accumulation, OxLDL uptake and the formation of OxLDL-induced MΦ foam cells. Additionally, we show that SPA deficiency markedly attenuated AS development. To our knowledge, this study is the first to report direct evidence of the pro-atherogenic effects of SPA.

Foam cells play a major role in plaque formation in AS^23^, and our study provides clear evidence about the involvement of SPA in MΦ-derived foam cells in early atherosclerosis. The formation and retention of MΦ foam cells in lesions exacerbate the disease and the development of vulnerable plaque^32^. Thus, strategies targeting the formation of foam cells can slow lesion progression and may serve as a potential adjunctive treatment for lipid-decreasing therapies^32^. SPA appears to be a positive regulator controlling MFCF, and the upregulation of SPA plays a causative role in AS development. Thus, targeting SPA may be a promising therapeutic strategy.

Lipid accumulation is essential for macrophage differentiation and foam cell formation. The scavenger receptor CD36 is expressed on the cell surface of macrophages and has been found to be integral in the lipid-uptake process^26,33^. We found that SPA deficiency inhibits CD36 cellular expression with a corresponding decrease in CD36 mRNA levels. Our findings are consistent with a previous report showing that SPA can redistribute CD36 to the cell membrane of alveolar MΦs and also promotes CD36 expression, which facilitates surfactant lipid uptake by the alveolar MΦs^34^. Our findings support that SPA mediates OxLDL uptake and MFCF via CD36.

One limitation to this study is the use of SPA global knockout mice, which does not address the cell-specific role of SPA in AS. In addition to macrophages, smooth muscle and adventitia cells could also express SPA. The cell-specific roles of SPA in AS may be investigated in the future by using tissue-specific SPA deficient mice.

In summary, this study has demonstrated a novel role for SPA in promoting the development of MΦ foam cell formation and atherosclerosis. These findings suggest that the modulation of SPA could be a novel approach for reducing atherosclerosis.

## ACKNOWLEDGMENTS

We would like to thank Dr. Lu Hong for providing *hPCSK9 (PCSK9^DY^*) AAV^20^.

## Supplemental Materials

Detailed Methods

Online Table I

Major Resources Table

## HIGHLIGHTS

- Surfactant protein A (SPA) is a novel protein regulating macrophage-derived foam cell formation.
- SPA modulates macrophage foam cell formation by regulating CD36 expression.
- SPA deficiency attenuates the early development of atherosclerosis.

## REFERENCES

1. Vasan RS, Benjamin EJ. The future of cardiovascular epidemiology. Circulation. 2016;133(25). doi:10.1161/CIRCULATIONAHA.116.023528

2. Heart Disease and Stroke Statistics-2021 Update A Report from the American Heart Association. Circulation. Published online 2021.doi:10.1161/CIR.0000000000000950

3. Hansson GKMDP. MECHANISMS OF DISEASE: Inflammation, Atherosclerosis, and Coronary Artery Disease. N Engl J Med. 2005;352(16).

4. Moore KJ, Sheedy FJ, Fisher EA. Macrophages in atherosclerosis: A dynamic balance. Nat Rev Immunol. 2013;13(10). doi:10.1038/nri3520

5. Chistiakov DA, Bobryshev Y V., Orekhov AN. Macrophage-mediated cholesterol handling in atherosclerosis. J Cell Mol Med. 2016;20(1). doi:10.1111/jcmm.12689

6. Yuan Y, Li P, Ye J. Lipid homeostasis and the formation of macrophage-derived foam cells in atherosclerosis. Protein Cell. 2012;3(3).doi:10.1007/s13238-012-2025-6

7. King SD, Chen SY. Recent progress on surfactant protein A: Cellular function in lung and kidney disease development. Am J Physiol - Cell Physiol. 2020;319(2):C316–C320. doi:10.1152/ajpcell.00195.2020

8. Ran R, Cai D, King SD, Que X, Bath JM, Chen Sy. Surfactant Protein A, a Novel Regulator for Smooth Muscle Phenotypic Modulation and Vascular Remodeling—Brief Report. Arterioscler Thromb Vasc Biol. 2021;41(2). doi:10.1161/atvbaha.120.314622

9. McNeely TB, Coonrod JD. Comparison of the opsonic activity of human surfactant protein a for staphylococcus aureus and streptococcus pneumoniae with rabbit and human macrophages. J Infect Dis. 1993;167(1). doi:10.1093/infdis/167.1.91

10. McNeely TB, Coonrod JD. Aggregation and opsonization of type A but not type B Hemophilus influenzae by surfactant protein A. Am J Respir Cell Mol Biol. 1994;11(1). doi:10.1165/ajrcmb.11.1.8018334

11. Wright JR. Immunomodulatory functions of surfactant. Physiol Rev. Published online 1997. doi:10.1152/physrev.1997.77.4.931

12. Weissbach S, Neuendank A, Pettersson M, Schaberg T, Pison U. Surfactant protein A modulates release of reactive oxygen species from alveolar macrophages. Am J Physiol - Lung Cell Mol Physiol. 1994;267(611-6). doi:10.1152/ajplung.1994.267.6.l660

13. Nozaki S, Kashiwagi H, Yamashita S, Nakagawa T, Kostner B, Tomiyama Y, Nakata A, Ishigami M, Miyagawa JI, Kameda-Takemura K, Kurata Y, Matsuzawa Y. Reduced uptake of oxidized low density lipoproteins in monocyte-derived macrophages from CD36-deficient subjects. J Clin Invest. 1995;96(4). doi:10.1172/JCI118231

14. Zhang X, Goncalves R, Mosser DM. The isolation and characterization of murine macrophages. Curr Protoc Immunol. 2008;(SUPPL. 83). doi:10.1002/0471142735.im1401s83

15. Leenaars M, Hendriksen CFM. Critical steps in the production of polyclonal and monoclonal antibodies: Evaluation and recommendations. ILAR J.2005;46(3). doi:10.1093/ilar.46.3.269

16. McCloy RA, Rogers S, Caldon CE, Lorca T, Castro A, Burgess A. Partial inhibition of Cdk1 in G2 phase overrides the SAC and decouples mitotic events. Cell Cycle. 2014;13(9). doi:10.4161/cc.28401

17. Toni LS, Garcia AM, Jeffrey DA, Jiang X, Stauffer BL, Miyamoto SD, Sucharov CC. Optimization of phenol-chloroform RNA extraction. MethodsX. 2018;5:599-608. doi:10.1016/j.mex.2018.05.011

18. García-Vence M, Chantada-Vazquez M del P, Sosa-Fajardo A, Agra R, Barcia de la Iglesia A, Otero-Glez A, García-González M, Cameselle-Teijeiro JM, Nuñez C, Bravo JJ, Bravo SB. Protein Extraction From FFPE Kidney Tissue Samples: A Review of the Literature and Characterization of Techniques. Front Med. 2021;8. doi:10.3389/fmed.2021.657313

19. Wang Y, Dubland JA, Allahverdian S, Asonye E, Sahin B, Jaw JE, Sin DD, Seidman MA, Leeper NJ, Francis GA. Smooth Muscle Cells Contribute the Majority of Foam Cells in ApoE (Apolipoprotein E)-Deficient Mouse Atherosclerosis. Arterioscler Thromb Vasc Biol. 2019;39(5):876–887. doi:10.1161/ATVBAHA.119.312434

20. Bjørklund MM, Hollensen AK, Hagensen MK, Dagnæs-Hansen F, Christoffersen C, Mikkelsen JG, Bentzon JF. Induction of atherosclerosis in mice and hamsters without germline genetic engineering. Circ Res. 2014;114(11). doi:10.1161/CIRCRESAHA.114.302937

21. Marx N, Sukhova G, Murphy C, Libby P, Plutzky J. Macrophages in human atheroma contain PPARgamma: differentiation-dependent peroxisomal proliferator-activated receptor gamma(PPARgamma) expression and reduction of MMP-9 activity through PPARgamma activation in mononuclear phagocytes in vitro. Am J Pathol. 1998;153(1): 17–23. doi:10.1016/S0002-9440(10)65540-X

22. Ricote M, Huang J, Fajas L, Li A, Welch J, Najib J, Witztum JL, Auwerx J, Palinski W, Glass CK. Expression of the peroxisome proliferator-activated receptor y (PPARy) in human atherosclerosis and regulation in macrophages by colony stimulating factors and oxidized low density lipoprotein. Proc Natl Acad Sci U S A. 1998;95(13). doi:10.1073/pnas.95.13.7614

23. Tabas I. Macrophage death and defective inflammation resolution in atherosclerosis. Nat Rev Immunol. 2010;10(1). doi:10.1038/nri2675

24. Liu Z, Zhu H, Dai X, Wang C, Ding Y, Song P, Zou MH. Macrophage Liver Kinase B1 Inhibits Foam Cell Formation and Atherosclerosis. Circ Res.2017;121(9). doi:10.1161/CIRCRESAHA.117.311546

25. Dai XY, Cai Y, Sun W, Ding Y, Wang W, Kong W, Tang C, Zhu Y, Xu MJ, Wang X. Intermedin inhibits macrophage foam-cell formation via tristetraprolin-mediated decay of CD36 mRNA. Cardiovasc Res. 2014;101(2). doi:10.1093/cvr/cvt254

26. Kunjathoor V.V., Febbraio M, Podrez EA, Moore KJ, Andersson L, Koehn S, Rhee JS, Silverstein R, Hoff HF, Freeman MW. Scavenger receptors class A-I/II and CD36 are the principal receptors responsible for the uptake of modified low density lipoprotein leading to lipid loading in macrophages. J Biol Chem. 2002;277(51). doi:10.1074/jbc.M209649200

27. Febbraio M, Podrez EA, Smith JD, Hajjar DP, Hazen SL, Hoff HF, Sharma K, Silverstein RL. Targeted disruption of the class B, scavenger receptor CD36 protects against atherosclerotic lesion development in mice. J Clin Invest. 2000;105(8). doi:10.1172/JCI9259

28. Podrez EA, Febbraio M, Sheibani N, Schmitt D, Silverstein RL, Hajjar DP, Cohen PA, Frazier WA, Hoff HF, Hazen SL. Macrophage scavenger receptor CD36 is the major receptor for LDL modified by monocyte-generated reactive nitrogen species. J Clin Invest. 2000;105(8). doi:10.1172/JCI8574

29. Nagy L, Tontonoz P, Alvarez JGA, Chen H, Evans RM. Oxidized LDL regulates macrophage gene expression through ligand activation of PPARγ. Cell. 1998;93(2). doi:10.1016/S0092-8674(00)81574-3

30. Emini Veseli B, Perrotta P, De Meyer GRA, Roth L, Van der Donckt C, Martinet W, De Meyer GRY. Animal models of atherosclerosis. Eur J Pharmacol. 2017;816. doi:10.1016/j.ejphar.2017.05.010

31. Glass CK, Witztum JL. Atherosclerosis: The road ahead. Cell. 2001;104(4). doi:10.1016/S0092-8674(01)00238-0

32. Libby P, Tabas I, Fredman G, Fisher EA. Inflammation and its resolution as determinants of acute coronary syndromes. Circ Res. 2014;114(12). doi:10.1161/CIRCRESAHA.114.302699

33. Rahaman SO, Lennon DJ, Febbraio M, Podrez EA, Hazen SL, Silverstein RLL. A CD36-dependent signaling cascade is necessary for macrophage foam cell formation. Cell Metab. 2006;4(3). doi:10.1016/j.cmet.2006.06.007

34. Dodd CE, Pyle CJ, Glowinski R, Rajaram MVS, Schlesinger LS. CD36-Mediated Uptake of Surfactant Lipids by Human Macrophages Promotes Intracellular Growth of Mycobacterium tuberculosis. J Immunol. 2016;197(12). doi:10.4049/jimmunol.1600856

