## Supplementary Materials for "Surfactant protein A promotes atherosclerosis through mediating macrophage foam cell formation"

#### Detailed Methods

##### *Expression and purification of SPA protein*

The purification of the SPA protein was done as previously described[8]. Briefly, human SPA cDNA clone in the pDONR221 (DNASU clone# HsCD00951577) was recombined into the pDEST17 gateway destination vector using LR clonase II enzyme mix. The pDEST-SPA vector was transformed into BL21(DE3) competent cells, and SPA protein expression was induced with 1 mmol/L isopropyl β-d-thiogalactopyranoside (IPTG) for 24 h. SPA proteins were extracted and purified using the HisTrap HP system according to the manufacturer's instructions. The SPA protein was concentrated using Microcon-10 kDa centrifugal filter unit.

##### *SPA antibody production*

The SPA polyclonal antibody was produced by using the purified SPA recombinant proteins as the antigen which was subcutaneously injected with Freunds Complete Adjuvant into 4 injection sites of ten 6-week-old mice as described[15]. Two weeks later, the same dosage was administrated for booster purpose. The antibody response was monitored by obtaining and evaluating a blood sample for antibodies in the serum. Exsanguination was performed 4 weeks later under general anesthesia via cardiac puncture. Antibodies were collected from serum and purified by liquid chromatography via affinity selection using the SPA protein.

##### *SPA immunofluorescent staining in vitro*

Isolated macrophages were cultured on coverslips placed in a 24 well plate. Cells were seeded on coverslips in a 24 well plate and cultured in supplemented DMEM. The next day, cells were rinsed with PBS then fixed in 10% formalin for 10 minutes at room temperature (RT) then blocked with 10% goat serum in PBS-T for 30 minutes at RT then incubated with SPA primary antibody (1:100) for 1 h at RT. After primary antibody incubation cells were rinsed twice with PBS-T and incubated with the corresponding secondary antibody (1:200) for 30 minutes. Coverslips were rinsed twice with PBS-T and mounted on glass slides with Prolong Gold antifade mounting medium containing DAPI. Fluorescent images were captured using a Keyence microscope and processed using ImageJ.

##### *BODIPY staining*

Isolated macrophages were cultured on coverslips placed in a 24 well plate in DMEM without FBS and treated with vehicle (control) or OxLDL (50 µg/ml) for 24 h. Cells were rinsed with PBS and fixed in 10% formalin for 10 minutes at room temperature (RT), cells were then washed three times with PBS-T and incubated with 10µg/ml BODIPY staining solution in the dark at RT for 30 minutes. Samples were washed with PBS-T three times and mounted on glass slides with Prolong Gold antifade mounting medium containing DAPI. Fluorescent images were captured using a Keyence microscope and processed using ImageJ. The fluorescence intensity per cell was calculated with ImageJ using the following formula to calculate the Corrected Total Cell Fluorescence, CTCF= Integrated Density-(area of selected cell* Mean background fluorescence), as previously described[16].

***Filipin III Staining***

The cell membrane cholesterol levels were detected using Filipin III staining. Isolated macrophages were cultured on coverslips placed in a 24 well plate in DMEM without FBS and treated with vehicle (control) or OxLDL (50 µg/ml) for 24 h. After rinsing cells with PBS, cells were fixed in 10% formalin for 10 minutes at RT. Cells were rinsed twice with Tris Buffer and incubated with Filipin III stock solution diluted at 1:100 in Tris Buffer for 30 minutes in the dark. Coverslips were washed 2 times for 5 min each with Tris Buffer and then mounted with glycerol followed by immediate observation of cell staining using a Keyence microscope. The fluorescence intensity per cell was calculated with ImageJ using the following formula to calculate the Corrected Total Cell Fluorescence, CTCF= Integrated Density-(area of selected cell* Mean background fluorescence), as previously described[16].

***DiI-labeled Ox-LDL uptake and macrophage foam cell formation assay***

OxLDL uptake and foam cell formation were analyzed using DiI-labeled OxLDL. Isolated macrophages were cultured on coverslips placed in a 24 well plate in DMEM without FBS and treated with vehicle (control) or OxLDL (50 µg/ml) for 24 h. Cells were washed with PBS then fixed in 10% formalin for 10 minutes, rinsed with PBS three times, then mounted onto glass slides with Prolong Gold antifade mounting medium containing DAPI. Fluorescent images were captured using a Keyence microscope and processed using ImageJ. The % of foam cells was calculated by counting the number of cells containing DiI-OxLDL uptake and dividing it by the total number of cells. The fluorescence intensity per cell was calculated with ImageJ using the following formula to calculate the Corrected Total Cell Fluorescence, CTCF= Integrated Density-(area of selected cell* Mean background fluorescence), as previously described[16].

***Cellular expression of CD36***

To detect the cellular expression of CD36, primary mouse peritoneal resident macrophages were treated with serum free DMEM media with the vehicle (control) or OxLDL (50 µg/ml) treatment for 24 h. Treated cells were rinsed first with PBS then fixed with ice cold methanol for 15 minutes at 4° C and subsequently rinsed 3 times with PBS. Coverslips were incubated in 10% goat serum in PBS-T at RT for 30 minutes then incubated with primary antibodies CD36 (1:100) for 1 h, followed by two washes with PBS-T then incubated with FITC-labeled secondary antibody for 30 minutes. Samples were washed 3 times with PBS-T and mounted on glass slides with Prolong Gold antifade mounting medium containing DAPI. Fluorescent images were captured using a Keyence microscope and processed using ImageJ. The fluorescence intensity per cell was calculated with ImageJ using the formula to calculate the Corrected Total Cell Fluorescence, CTCF= Integrated Density-(area of selected cell* Mean background fluorescence), as previously described[16].

***RNA extraction and real-time quantitative PCR***

Cells were collected, and total RNA was extracted using Trizol Reagent as previously described[17]. 1 µg of total RNA was reverse transcribed using RevertAid First Strand cDNA Synthesis Kit according to the manufacturers protocol. qPCR was performed in triplicates using All-in-One qPCR Mix using a Mx3005P qPCR System (Agilent Technologies). Primer sequences are included in the Major Resources Table.

***Adeno-associated virus (AAV) administration***

The pAAV/D374Y-*hPCSK9* (*PCSK9*$^{DY}$) plasmid driven by the ApoEHCR-hAAT promoter[20] was a gift from Dr. Lu Hong (Addgene plasmid # 58379). The *AAV-PCSK9*$^{DY}$ virus was delivered in a single tail-vein injection as previously described[20]. One week post injection, animals were fed a high fat diet for 4 or 6 weeks as indicated to induce AS.

***Western blot analysis***

Tissues were collected in RIPA lysis buffer containing phosphatase and protease inhibitors, and protein extraction was conducted as previously described[8]. Subsequently, equal amounts of proteins were separated on 8-12% SDS-PAGE and transferred to nitrocellulose membrane. Membranes were blocked with 5% BSA in TBS-T for 1 h and probed with primary antibodies against the proteins of interest followed by incubation with secondary antibodies. Protein bands were detected and analyzed using an Odyssey CLx Imager imaging system and processed using Image Studio Software (ver 5.2 software; Li-Cor Biosciences).

***Brachiocephalic artery histological and immunofluorescent staining***

For the analysis of SPA expression in early AS lesions, wild-type (WT) and ApoE$^{-/-}$ mice were fed a high fat diet for 4 weeks. Mouse brachiocephalic arteries (BCAs) were fixed in 4% paraformaldehyde overnight and then dehydrated in a series of ethanol and xylene solutions, embedded in paraffin and sectioned in 5µm sections. Next, hydrated samples underwent antigen retrieval and blocking then were incubated with SPA (1:100) or F4/80 (1:100) primary antibody or corresponding normal IgG overnight at 4$^{o}$C. The next day, slides were incubated with FITC or TRITC-labeled secondary antibodies (1:200) at RT for

30 min, rinsed twice with PBS and mounted with a coverslip containing Prolong Gold antifade mounting medium with DAPI. Fluorescent images were captured using a Keyence microscope. Images were processed using ImageJ software.

To analyze AS lesions, mouse BCA were fixed in 4% paraformaldehyde overnight and then dehydrated in a serious of ethanol and xylene solutions, embedded in paraffin and sectioned in 5 μm sections. The sections were deparaffinized in xylene and rehydrated and stained with hematoxylin (H&E) to assess for AS lesion development.

For the analysis of macrophage foam cells *in vivo*, serial sections (5 μm) of OCT-embedded frozen BCA tissues were washed 3 times with PBS for 5 min then blocked with 10% goat serum for 30 min and stained simultaneously for (a) F4/80 (1:100) and (b) cholesteryl ester (BODIPY 493/503; 10 μg/ml). F4/80 (1:200) primary antibody was incubated overnight at 4°C as described[19]. The next day, slides were incubated with TRITC-labeled secondary antibodies (1:200) containing BODIPY at room temperature for 30min[4], rinsed twice and mounted with a coverslip containing Prolong Gold antifade mounting medium with DAPI. Sections stained with BODIPY and IgG specific secondary served negative controls. Only BODIPY+ areas that colocalized with DNA were considered to be foam cells; BODIPY+ areas that did not colocalize with DNA were designated as extracellular deposits of cholesteryl ester and were not counted as foam cells. Colocalization of red (F4/80) with green (foam cells) were scored as containing yellow or not and counted as macrophage foam cells if yellow was observed as previously described[19]. Images were captured using a Keyence microscope and were processed using ImageJ.

**Online Table I: Sample size, normality tests, statistical tests and p values for data presented in all Figures.**

| Figure | Group | Size | Normality Test | Pass | Statistical test | p value |
|---|---|---|---|---|---|---|
| Fig. 1C | WT Ctrl | 49 | Shapiro-Wilk test | No | Krustal-Wallis test followed by Dunn's multiple comparisons test | *: WT Ctrl vs. WT OxLDL p< 0.0001; WT Ctrl vs. SPA-/- Control p= 0.0003 |
| | WT OxLDL | 55 | | | | #: < 0.0001 |
| | SPA-/- Ctrl | 45 | | | | $: < 0.0001 |
| | SPA-/- OxLDL | 45 | | | | |
| Fig. 1E | WT Ctrl | 18 | D'Agostino & Pearson test | Yes | Ordinary one-way ANOVA followed by Tukey's multiple comparisons test | *: WT Ctrl vs. WT OxLDL p< 0.0001; WT Ctrl vs. SPA-/- OxLDL p= 0.034 |
| | WT OxLDL | 31 | | | | #: <0.0001 |
| | SPA-/- Ctrl | 23 | | | | $: 0.031 |
| | SPA-/- OxLDL | 31 | | | | |
| Fig. 1G | WT OxLDL | 36 | Shapiro-Wilk test | No | Mann-Whitney test | *: <0.0001 |

| | | | | | | |
|---|---|---|---|---|---|---|
| | SPA-/- OxLDL | 40 | | | | |
| Fig. 1H | WT OxLDL | 9 | Shapiro-Wilk test | Yes | Unpaired two-tailed t test with Welch's correction | *: <0.0001 |
| | SPA-/- OxLDL | 9 | | | | |
| Fig. 1J | WT Ctrl | 60 | Shapiro-Wilk test | No | Kruskal-Wallis test followed by Dunn's multiple comparisons test | *: p<0.0001 |
| | WT OxLDL | 67 | | | | #: WT OxLDL vs. SPA-/- Ctrl p<0.0001; WT OxLDL vs SPA-/- OxLDL p= 0.046 |
| | SPA-/- Ctrl | 62 | | | | $: < 0.0001 |
| | SPA-/- OxLDL | 88 | | | | |
| Fig. 1K | WT Ctrl | 7 | Shapiro-Wilk test | Yes | Ordinary one-way ANOVA followed by Tukey's multiple comparisons test | *: WT Ctrl vs. WT OxLDL p <0.0001; WT Ctrl vs. SPA-/- Ctrl p=0.032; WT Ctrl vs. SPA-/- OxLDL p= 0.002 |
| | WT OxLDL | 7 | | | | #: <0.0001 |
| | SPA-/- Ctrl | 8 | | | | |
| | SPA-/- OxLDL | 8 | | | | |
| Fig. 2B | Ctrl | 3 | N/A | | Nonparametric two-tailed Mann-Whitney test | 0.100 |
| | AS | 3 | | | | |
| Fig. 2E | Ctrl | 3 | N/A | | Nonparametric two-tailed Mann-Whitney test | *: 0.029 |
| | AS | 3 | | | | |
| Fig. 2H | WT | 4 | N/A | | Nonparametric two-tailed Mann-Whitney test | *: 0.029 |
| | SPA-/- | 4 | | | | |
| Fig. 2J | WT | 5 | N/A | | Nonparametric two-tailed Mann-Whitney test | *: 0.008 |
| | SPA-/- | 5 | | | | |

### Major Resources Table

#### Animals (*in vivo* studies)

| Species | Vendor/Source | Background strain | Sex | Persistent ID/URL |
|---|---|---|---|---|
| *SPA* $^{-/-}$ (*Sftpa1* $^{tm1kor}$) | Jackson Laboratory | BL6. 129P2 | M, F | Strain #: 004964 |
| ApoE $^{-/-}$ (*Apoe* $^{tm1Unc}$/J) | Jackson Laboratory | B6.129 | M, F | Strain #: 002052 |
| WT C57BL6/J | Jackson Laboratory | BL6 | M, F | Strain #: 000664 |

#### Genotyping primers 5'- 3'

##### Sftpa1$^{tm1kor}$

| Primer 1 | Primer 2 | Primer 3 |
|---|---|---|
| GCTACTTCCATTTGTCAC GTCC | ACAGAAGTTTGTGCCGGAA G | ATGGTCACCCAGAAAA CAGG |

##### Apoe$^{tm1Unc}$/J

| Primer 1 | Primer 2 | Primer 3 |
|---|---|---|
| TGTGACTTGGGAGCTCTG CAGC | GCCGCCCCGACTGCATCT | GCCTAGCCGAGGGAGAGCC G |

#### Antibodies

| Target Antigen: Host | Vendor/Source | Catalog # | Working Concentration | Persistent ID/URL |
|---|---|---|---|---|
| SPA: Mouse | This study | N/A | IF: (1:100) | N/A |
| CD68: Rabbit | ThermoFisher Scientific | PA5-78996 | IF: (1:200) | |
| F4/80: Rat | Abcam | Ab16911 | IF: (1:100) | |
| α Tubulin | Sigma Aldrich | T5168 | WB: (1:1,000) | |
| CD36 (D8L9T): Rabbit | Cell Signaling | mAb#14347 | IF: (1:200) | |
| Anti-rat IgG (H+L) (Alexa Fluor 555 Conjugate) | Cell Signaling | 4417 | IF (1:200) | |
| Goat anti-Rabbit IgG (H+L), FITC | Invitrogen | 31635 | IF (1:200) | Goat anti-Rabbit IgG (H+L), FITC (31635) (thermofisher.com) |
| Goat anti-Mouse IgG (H+L) Texas Red-X | Invitrogen | T-862 | IF (1:200) | Goat anti-Mouse IgG (H+L) Cross-Adsorbed, Texas Red-X (T-862) (thermofisher.com) |

| | | | | |
|---|---|---|---|---|
| Goat anti-mouse IgG (H+L) Alexa Fluor 488 | Invitrogen | R37120 | IF (1:200) | |
| Goat anti-rabbit IgG (H+L) Alexa Fluor 568 | Invitrogen | A-11011 | IF (1:200) | |
| IRDye 680RD Goat anti-mouse IgG | IRDye | 926-68071 | WB (1:10,000) | IRDye 680RD Goat anti-Mouse IgG Secondary Antibody (licor.com) |
| IRDye 680RD Goat anti-rabbit IgG | IRDye | 926-68071 | WB (1:10,000) | IRDye 680RD Goat anti-Rabbit IgG Secondary Antibody (licor.com) |

##### Cultured Cells

| Name | Vendor or Source | Sex | Persistent ID/URL |
|---|---|---|---|
| Mouse resident peritoneal macrophages | Primary culture Chen Lab | M, F | N/A |

##### DNA/cDNA Clones

| Clone Name | Plasmid ID | Source | Persistent ID/URL |
|---|---|---|---|
| pDONR221_SFTPA1 | Clone# HsCD00951577 | DNASU Plasmid Repository | https://dnasu.org/DNASU/AdvancedSearchOptions.do |
| pAAV/D374Y-hPCSK9 | Addgene #58379 | Gift from Dr. Lu Hong | *PCSK9*$^{DY}$ |

##### Primers for RT-PCR

| Name | Forward primer 5'-3' | Reverse primer 5'-3' |
|---|---|---|
| CD36 | TGAATGGTTGAGACCCCGTG | TACGTGGCCCGGTTCTACTA |
| CYP | CAACTCCCTCAAGATTGTCAGCAA | GGCATGGACTGTGGTCATGA |

##### Reagents

| Name | Vendor or Source | Catalog number |
|---|---|---|
| L-glutamine | Hyclone | SH30034.01 |
| DMEM | Corning | 17-207-CV |
| FBS | R&D Systems | S11550H |
| OxLDL | ThermoFisher Scientific | L34357 |

| | | |
|---|---|---|
| Bodipy493/503 | ThermoFisher Scientific | D3922 |
| Filipin III | Cayman Chemical | NC9384264 |
| Dil- OxLDL | ThermoFisher Scientific | L34358 |
| Prolong Gold Antifade Mountant with DAPI | ThermoFisher Scientific | P36931 |
| Protease inhibitor | Sigma Aldrich | P8340 |
| Phosphatase Inhibitor | Sigma Aldrich | P0044 |
| 10% normal goat serum | Fisher Scientific | 50-675-69 |
| Trizol | ThermoFisher Scientific | 10296028 |
| All-in-One qPCR Mix | GeneCopoeia | QP001 |
| Hematoxylin and eosin kit | American Master Tech | KTHNEPT |
| isopropyl β-d-thiogalactopyranoside | ThermoFisher Scientific | 15529019 |
| xTractor Buffer | Takara, Japan | 63562 |
| Capturem His-tagged purification miniprep kit | Takara, Japan | 635710 |
| Microcon-10 kDa centrifugal filter unit | Millipore | MRCPRT010 |
| RevertAid First Strand cDNA Synthesis Kit | ThermoFisher Scientific | K1621 |
| Recombinant SPA protein | This study | N/A |
| Gateway pDEST17 Vector | ThermoFisher Scientific | 11803012 |
| LR Clonase II Plus enzyme | ThermoFisher Scientific | 12538120 |
| BL21(DE3) Competent *E. coli* | New England Biolabs | C2527H |
| HisTrap HP | GE Healthcare | 71-5027-68AF |
| Freunds Complete Adjuvant | Thermofisher | 77140 |
| Pierce Antibody Clean-up Kit | Thermofisher | 44600 |
